# MtbEis protein: A new player in stringent response of *Mycobacterium tuberculosis*

**DOI:** 10.1101/2023.10.29.564617

**Authors:** Nittu Singh, Vandana Basra, Charu Sharma

## Abstract

The stringent response in bacteria is an important defense mechanism in which hyperphosphorylated forms of guanosine, also known as molecular alarmones, are synthesized by RelA/SpoT Homolog (RSH) proteins. Rel protein in *Mycobacterium tuberculosis* (Rel_Mtb_) also regulates expression of persistence or virulence associated genes. Loss of Rel_Mtb_ leads to higher expression of few of the virulence and cell wall remodeling factors in addition to upregulating many secreted antigens and proteins. The <u>E</u>nhanced Intracellular <u>S</u>urvival (MtbEis) protein is one of the upregulated virulence factors. Based on this information, the precise role of MtbEis, a GNAT family acetyltransferase, in the stringent response in *M. tuberculosis* was explored. To begin with, MtbEis has been confirmed to enhance the guanosine pentaphosphate (pppGpp) synthesis activity of Rel_Mtb_ by acetylating Rel_Mtb_ at K513. Next, the *MtbEis* gene was knocked out in Mtb. The deletion of MtbEis resulted in a compromised survival accompanied by elevated levels of rRNAs under starvation conditions. Furthermore, the reduced expression of Rel_Mtb_ and subsequent decrease in pppGpp synthesis was also observed in MtbΔEis cells. Complementation of MtbΔEis with full length MtbEis restored the expression of Rel_Mtb_, rRNAs, pppGpp levels and survival of *M. tuberculosis*. However, MtbEis lacking acetyltransferase domain (ΔAT) failed to restore this, confirming the role of MtbEis mediated acetylation in regulating stringent response. In sum, our findings not only report the unexplored role of MtbEis in the starvation survival but also the acetylation of Rel_Mtb_ as a novel mechanistic aspect of stringent response in *M. tuberculosis*. In addition, this being the first post translational modification (PTM) report on any of the bacterial Rel proteins opens up the field for the discovery of new PTMs of Rel proteins.

**Author Summary:** Stringent Response is essential for bacterial survival and pathogenesis under stress conditions. The bifunctional Rel_Mtb_ enzyme is the key player of stringent response in *M. tuberculosis* catalyzing the synthesis of pppGpp from ATP and GTP under nutrient deprived conditions. Under stringent conditions, the tight regulation of transcriptional and translational processes aids the intracellular survival of Mtb. Amongst the translational regulations, the post-translational modification (PTM) is one of the most efficient mechanisms regulating the functions of enzymes. None of the PTMs of Rel proteins is known so far. Here, we have not only identified the acetylation of Rel_Mtb_ by MtbEis but also dissected the consequences of this modification in regulation of stringent response. MtbEis is a known virulence factor belonging to GNAT family acetyltransferase that is upregulated upon stringent response activation in Mtb. The MtbEis regulates expression of rRNAs and Rel_Mtb_, synthesis of pppGpp and Mtb survival by acetylating Rel_Mtb_ under starvation conditions. The regulation of stringent response by MtbEis mediated acetylation of Rel_Mtb_ have unravelled the novel function of MtbEis in adapting Mtb to existing environmental challenges associated with nutrient deficit state.

**Highlights:**

- None of the PTMs of Rel proteins is known so far
- MtbEis interacts with Rel_Mtb_ and acetylates Rel_Mtb_ at K513
- The acetyltransferase domain of MtbEis regulates rRNAs and Rel_Mtb_ expression, pppGpp synthesis and Mtb survival under starved conditions
- MtbEis helps Mtb in adapting to nutrient deficit state

## Introduction

Stringent response in bacterial kingdom is a conserved adaptive response that gets activated under different kinds of stresses. This response regulates global transcription to modify metabolism so that bacteria endures variety of stresses for a long duration (1). A hallmark feature of stringent response is the Rel protein mediated synthesis of hyperphosphorylated guanine nucleotides (pppGpp and ppGpp), commonly represented as (p)ppGpp. These hyperpohsphorylated molecules act as alarmones. After attaining sufficient levels, this signaling molecule in turn binds to the RNA polymerase and acts as negative regulator by decreasing the biosynthesis of stable RNAs including rRNA and tRNA and as a positive regulator by increasing the levels of mRNA transcripts of enzymes involved in amino acid biosynthesis and synthesis of proteins required for defense in stress (2–5).

The stringent response of *Mycobacterium tuberculosis* (Mtb) enables it to survive intracellularly under nutrient limiting conditions by synthesizing (p)ppGpp molecules. The synthesis and hydrolysis of (p)ppGpp in Mtb are mediated by synthetase and hydrolytic activity of bifunctional Rel_Mtb_ enzyme, respectively (6). The accumulation of (p)ppGpp and strict regulation of rRNAs are also conserved in stringent response of mycobacteria (7–9). The loss of Rel_Mtb_ leads to impaired survival of mycobacteria in growing cultures (7), animal models including mice (10) and guinea pigs (11). The global transcriptional study on Rel_Mtb_ deletion mutant (*Δrel*_Mtb_) revealed the modulation of expression of several genes in Mtb (10). Out of which, the inverse regulations of expression of only MtbEis and HspX by Rel_Mtb_ under stringent conditions are confirmed at protein levels (8).

MtbEis protein, being a member of GCN5-related family of N-acetyltransferases, acetylates aminoglycosides and proteins (12). The inactivation of aminoglycosides due to acetylation by MtbEis imparts drug resistance to mycobacteria (13). The MtbEis mediated acetylation at Lys55 of dual-specificity protein phosphatase 16/Mitogen activated protein kinase phosphatase-7 (DUSP16/MKP-7) protein of host led to its enhanced phosphatase activity resulting into concomitant inhibition of c-JNK signaling pathway and suppression of protective host immune response (14). MtbEis also acetylates histone H3 protein in IL-10 promoter which increases the expression of IL-10 and thereby suppressing autophagy (15). Besides, MtbEis mediated acetylation of nucleoid associated protein of Mtb (MtHU/HupB/Rv2986c) alters its DNA binding capacity leading to genome reorganization (16,17).

There is plenty of information available on MtbEis mediated acetylation of aminoglycosides and various proteins. However, based on the regulation of expression of MtbEis by Rel_Mtb_, we decided to investigate the unidentified role of MtbEis in stringent response by checking the interaction of Rel_Mtb_ with MtbEis and the outcome of these interactions during stringent conditions.

## RESULTS

### Rel_Mtb_ protein interacts with MtbEis

To examine the role of MtbEis in stringent response, the study was initiated by exploring the interaction between Rel_Mtb_ and MtbEis using *in vitro* GST pull down assay. The recombinant His-MtbEis, GST and GST-Rel_Mtb_ proteins purified to homogeneity were used (Fig. S1 Panels A, B and C). Only GST and GST-Rel_Mtb_ proteins were precipitated by GST-agarose beads ruling out the nonspecific association of His-MtbEis to either glutathione-agarose beads or GST proteins (Fig. 1A, Lanes 1-4). His-MtbEis protein was pulled down specifically by GST-Rel_Mtb_ confirming the interaction between His-MtbEis and GST-Rel_Mtb_ (Fig. 1A, Lane 5).

**Fig. 1:**
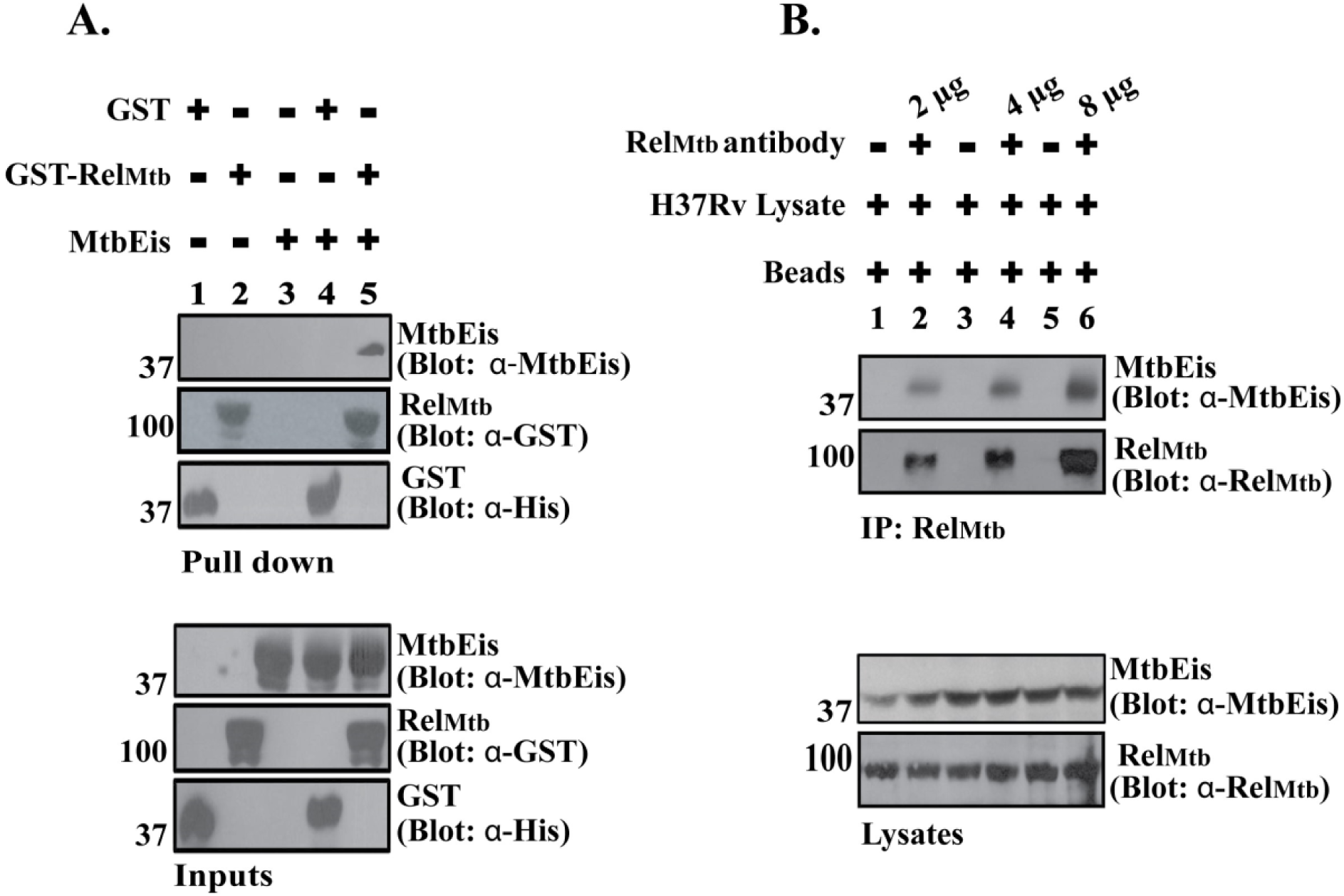
Rel_Mtb_ is a novel interacting partner of MtbEis: **Panel A.** *In vitro* GST Pull-Down assay was perfromed using GST-Rel_Mtb_, His-MtbEis and GST proteins as mentioned. Detection of GST-Rel_Mtb_ and GST proteins was done using anti-GST and anti-His antibody, respectively, while MtbEis was detected using anti-MtbEis antibody. Inputs panel depict the level of proteins used in the assay. **Panel B.** Immunoprecipitation was performed from lysates of Mtb using varying concentrations of anti-Rel_Mtb_ antibody. The mixture of lysate with Protein A beads served as a negative control. Detection of Endogenous Rel_Mtb_ and MtbEis proteins was done using anti-Rel_Mtb_ and anti-MtbEis antibodies. The levels of respective proteins present in the lysates are indicated in the lysates panel. The proteins used in the assay are shown at the top of figure. Molecular weight markers are indicated on the left. The blots are representation of three independent experiments. **Figure 1-Source data 1** **Source data 1 files for Figure 1 include raw and labelled images for the western blots shown in Panels A and B**.

To establish the interaction between MtbEis and native Rel_Mtb_ at the physiological level, the H37Rv lysate was subjected to immunoprecipitation using Rel_Mtb_ antibody. The absence of non-specific association of MtbEis with Protein A beads and the increased presence of MtbEis protein in the immunoprecipitates of Rel_Mtb_ upon increasing the concentration of Rel_Mtb_ antibody confirms the interaction of Rel_Mtb_ with MtbEis *in vivo* (Fig.1B).

### MtbEis enhances the accumulation of (p)ppGpp under *in vitro* conditions

Rel_Mtb_ is a bifunctional enzyme having synthetase and hydrolase activities (6). To check the effect of interaction of Rel_Mtb_ with MtbEis on its (p)ppGpp synthesis activity, the enzymatic assay was set up with recombinant His-Rel_Mtb_ and His-MtbEis proteins either alone or in combinations and the levels of (p)ppGpp formation was qualitatively analysed by TLC through autoradiography. The enhanced accumulation of (p)ppGpp was observed in the presence of His-MtbEis (Fig 2, Lanes 3 vs 4).

### Purified recombinant MtbEis protein acetylates Rel_Mtb_ at K513 residue

To check if MtbEis acetylates Rel_Mtb_, His-MtbEis and His-Rel_Mtb_ proteins were purified to homogeneity (Fig.S1, Panel A and Fig.S2, Panel A) and used for *in vitro* acetyltransferase assay. The incorporation of ^14^C-Acetyl group in Rel_Mtb_ indicated that MtbEis acetylates Rel_Mtb_ (Fig.3A, Lane 3, Upper Panel).

For the identification of amino acids of Rel_Mtb_ undergoing MtbEis mediated acetylation, purified Rel_Mtb_ before and after acetylation was subjected to trypsin digestion and then tandem mass spectrometry connected to liquid chromatography (LC-MS/MS). The peptide ^500^KLENGEVVEVFTS<u>K</u>APNAGPSR^521^ with molecular mass of 897.5 Da exhibited a mass of 939.5 Da after acetylation. The mass difference of 42 Da corresponding to acetyl group was identified on lysine residue at 513^th^ amino acid position in Rel_Mtb_ as a site of MtbEis mediated acetylation (Fig. 3B).

**Fig. 2:**
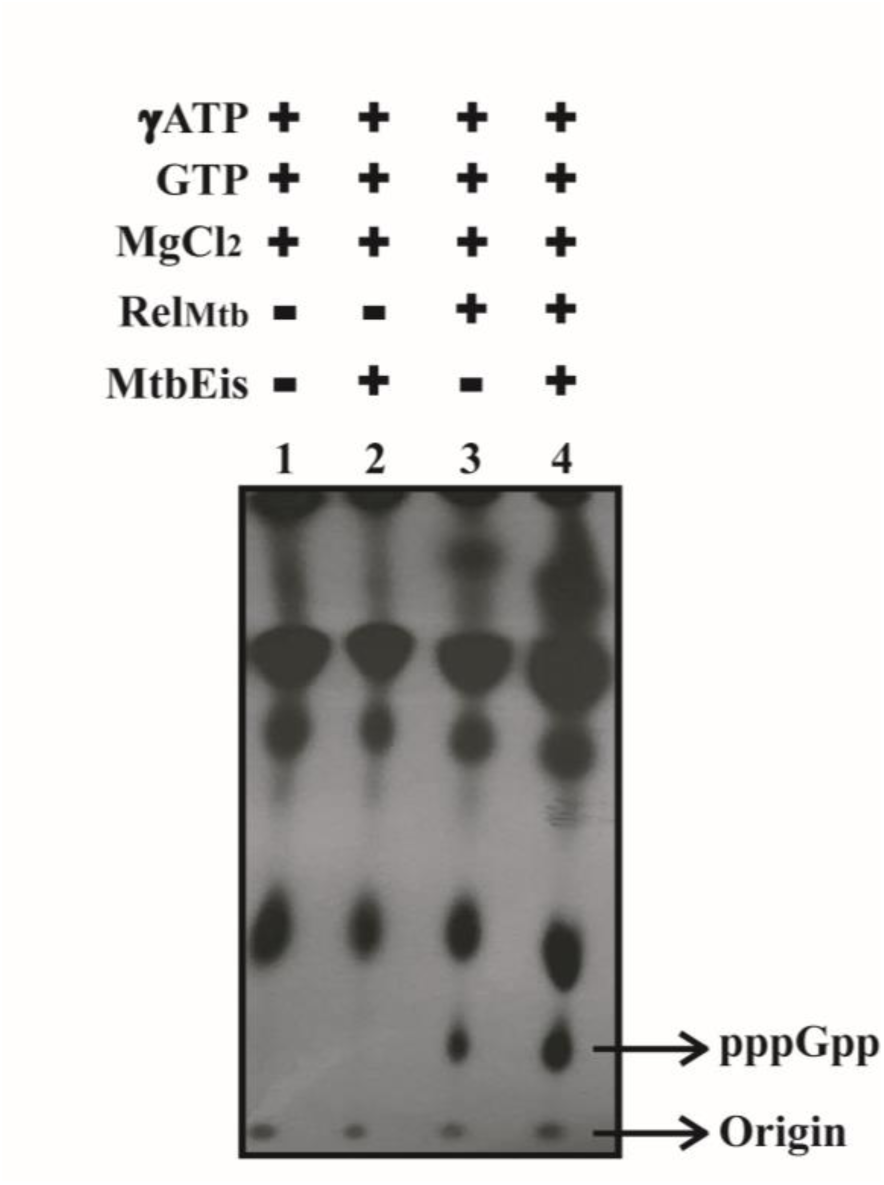
MtbEis enhances the accumulation of (p)ppGpp: (p)ppGpp synthesis by Rel_Mtb_ was carried out in the presence of MtbEis. The reaction mixture was resolved on PEI- TLC plates and analysed using autoradiography. The blot is a representation of three independent experiments. **Figure 2-Source data 1** **Source data files for Figure 2 include raw and labelled images for the blot shown in the figure**.

**Fig. 3:**
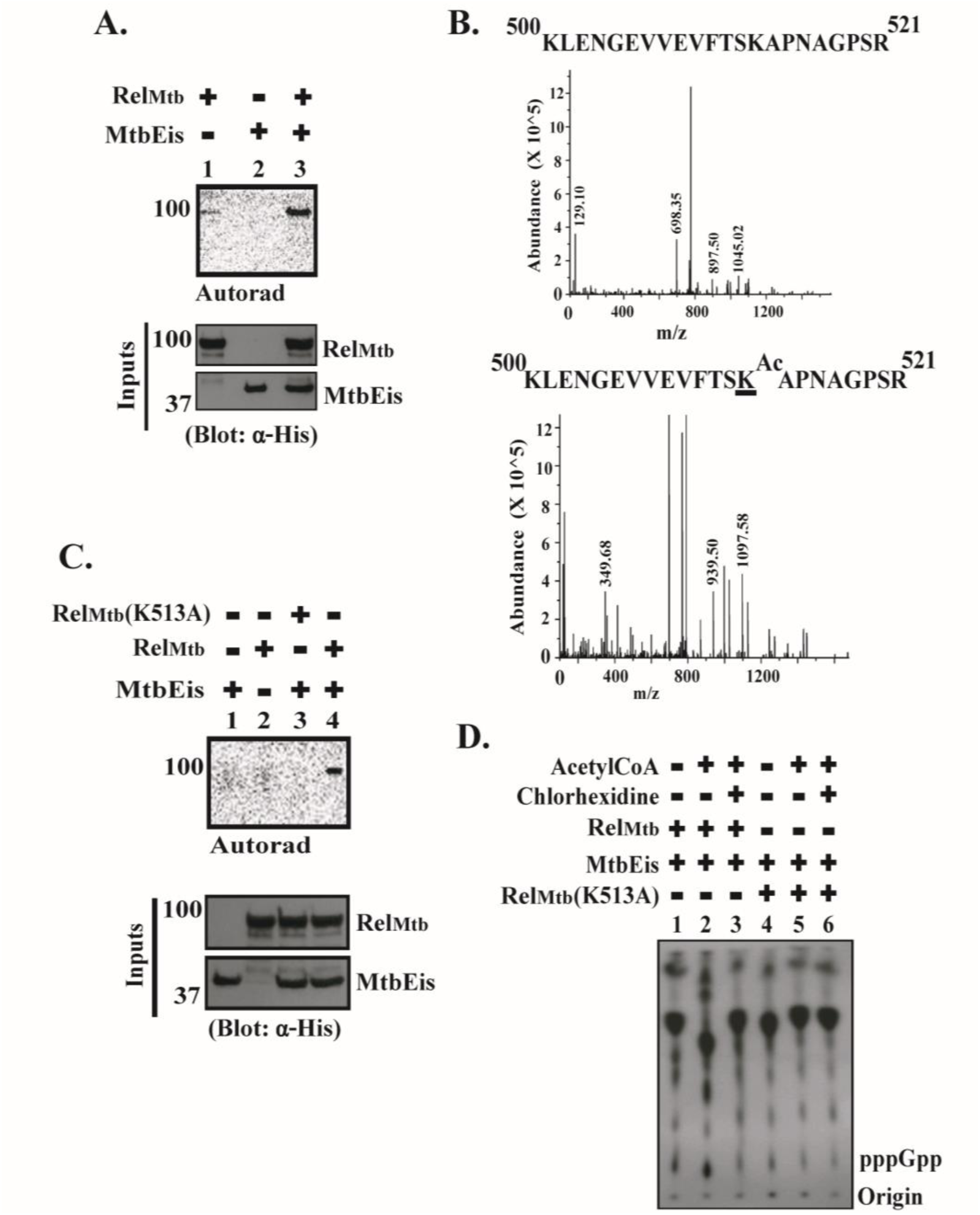
Rel_Mtb_ is acetylated by MtbEis at K513 residue: **Panel A.** *In-vitro* acetylation assay was set up with purified His-Rel_Mtb_ and His-MtbEis proteins using ^14^C-labelled acetyl-CoA as a substrate. The incorporation of ^14^C was analysed using autoradiography. Inputs indicate the level of proteins used for assay. Rel_Mtb_ and MtbEis were probed using anti-His antibody. **Panel B.** LC-MS/MS spectra of Rel_Mtb_ peptide 500-521 (^500^KLENGEVVEVFTSKAPNAGPSR^521^) after acetyltransferase reaction in the absence of MtbEis served as control (Upper Panel). LC-MS/MS spectra showing K513 as acetylation site in Rel_Mtb_ peptide (^500^KLENGEVVEVFTS**<u>K</u>^Ac^**APNAGPSR^521^) after acetyltransferase reaction in the presence of MtbEis (Lower Panel). The specific peak of acetylated lysine with m/z 939.50 Da showing the mass difference of 42 Da in peptide 500-521 is absent in spectra of control peptide. **Panel C.** The *in vitro* acetyltransferase reaction was performed using His-MtbEis, His-Rel_Mtb_ or His-Rel_Mtb_(K513A). The autorad shows the incorporation of ^14^C labelled Acetyl-CoA. The proteins used in this assay were probed using anti-His antibody and shown in input panels. **Panel D.** *In vitro* (p)ppGpp synthesis assay was performed as indicated above each lane. The reaction products were resolved on PEI–TLC plates and analysed using autoradiography. The molecular weight markers and names of the proteins in Panels A and C are indicated on left and right side. The blots and autorads are a representation of three independent experiments. **Figure 3-Source data 1** **Source data 1 files for Figure 3 include raw and labelled images for the blots shown in the Panels A, C and D**. **Figure 3-Source data 2** **Source data 2 files for Figure 3 include LC-MS/MS data for the images shown in the Panel B**.

To confirm that K513 is the site of MtbEis mediated acetylation of Rel_Mtb_ protein, the lysine residue at 513^th^ position was mutated to alanine. The wild type Rel_Mtb_ and its lysine mutant were purified to homogeneity (Fig. S2) and subjected to *in vitro* acetylation assay. The loss of incorporation of ^14^C-Acetyl group upon mutating lysine into Alanine of Rel_Mtb_ in comparison to wild type protein confirmed that Lys513 of Rel_Mtb_ is important for MtbEis mediated acetylation (Fig. 3C, Upper Panel, Lanes 3 vs 4).

### MtbEis mediated acetylation of Rel_Mtb_ at K513 residue enhances the accumulation of (p)ppGpp

The Acetyl-CoA is a well-known acetyl group donor (18). Therefore, to check the effect of MtbEis mediated acetylation on the synthesis activity of Rel_Mtb_, the *in vitro* (p)ppGpp synthesis assay was done in the presence of Acetyl-CoA. The enhanced accumulation of (p)ppGpp was observed in the presence of Acetyl-CoA (Fig. 3D, Lanes 1 vs 2). In order to further confirm whether accumulation of (p)ppGpp due to synthesis by Rel_Mtb_ is an acetylation dependent phenomenon, acetylation inhibitor, chlorhexidine, was used. The reduced levels of (p)ppGpp in the presence of chlorhexidine confirms the role of acetylation of Rel_Mtb_ in enhanced accumulation of (p)ppGpp (Fig. 3D, Lanes 2 vs 3). The significance of acetylation in enhanced accumulation of (p)ppGpp is further confirmed by only residual amount of (p)ppGpp in the presence of lysine mutant of His-Rel_Mtb_, i.e., Rel_Mtb_(K513A) (Fig. 3D, Lanes 1 and 2 vs Lanes 4 and 5). As expected, Rel_Mtb_(K513A), being acetylation defective mutant, did not respond to chlorhexidine (Fig. 3D, Lane 6).

### MtbEis mediated acetylation of Rel_Mtb_ influences the survival of *M. tuberculosis* and inhibits synthesis of rRNAs under starvation conditions

Cessation of growth rate is the physiological outcome of stringent conditions emanated due to synthesis of (p)ppGpp and halting of RNA synthesis (2,19). Therefore, the function of MtbEis in stringent response was validated by deleting MtbEis gene (*ΔMtbeis*) in H37Rv strain using recombineering method (Fig. S3, Panels A and B). The ΔMtbEis was complemented with full length MtbEis (ΔMtbEis::MtbEis) and MtbEis lacking acetyltransferase domain (ΔMtbEis::ΔAT) (Fig.S3, Panel C). The H37Rv strain lacking MtbEis and complemented strains were investigated for their survival ability under nutrients depleted conditions. At 0 h, the mycobacterial CFUs were similar for all the strains under both unstarved and starved conditions. Up to 24 h, the growth rate of ΔMtbEis or ΔAT strains was comparable to MtbH37Rv strain under unstarved conditions. However, under starved conditions, ΔMtbEis strain exhibited a greater loss in viability in comparison to wild type H37Rv. The fold difference between viability of H37Rv and ΔMtbEis strain was observed to be ∼4.0, ∼8.0 and ∼20.0 fold at 8.0 h, 16.0 h and 24.0 h of starvation, respectively. The loss in viability was restored when complemented with full length MtbEis but not by ΔAT (Fig.4, Panel A).

Next, the levels of accumulation of stable RNAs were determined under stringent conditions in H37Rv, MtbΔEis and complemented strains. The accumulation of 23S, 16S and 5S rRNA was higher by ∼13.0, ∼3.0 and ∼6.0 fold in MtbΔEis strain as compared to H37Rv strain. The accumulation of rRNAs in MtbΔEis strain was reduced to the levels of H37Rv strain on complementation with full length MtbEis but not with ΔAT (Fig.4, Panel B).

### Rel_Mtb_ protein expression is dependent on MtbEis mediated acetylation of Rel_Mtb_ under starvation conditions

Reduced survival with subsequent increase in rRNAs in MtbΔEis strain upon induction of stringent response implies that Rel_Mtb_ mediated functions are modulated by MtbEis. Therefore, the expression of Rel_Mtb_ in Wild type, MtbΔEis and complemented strains were checked after starvation of 24.0 h. The downregulation of Rel_Mtb_ protein in MtbΔEis strain by 3.5-fold in comparison to the wild type strain was observed (Fig. 5. Lane 2 vs. Lane 4). The Rel_Mtb_ protein levels were restored nearby normal levels only in MtbΔEis strain complemented with full length MtbEis but not with ΔAT (Fig. 5. Lanes 6 and 8).

**Fig. 4:**
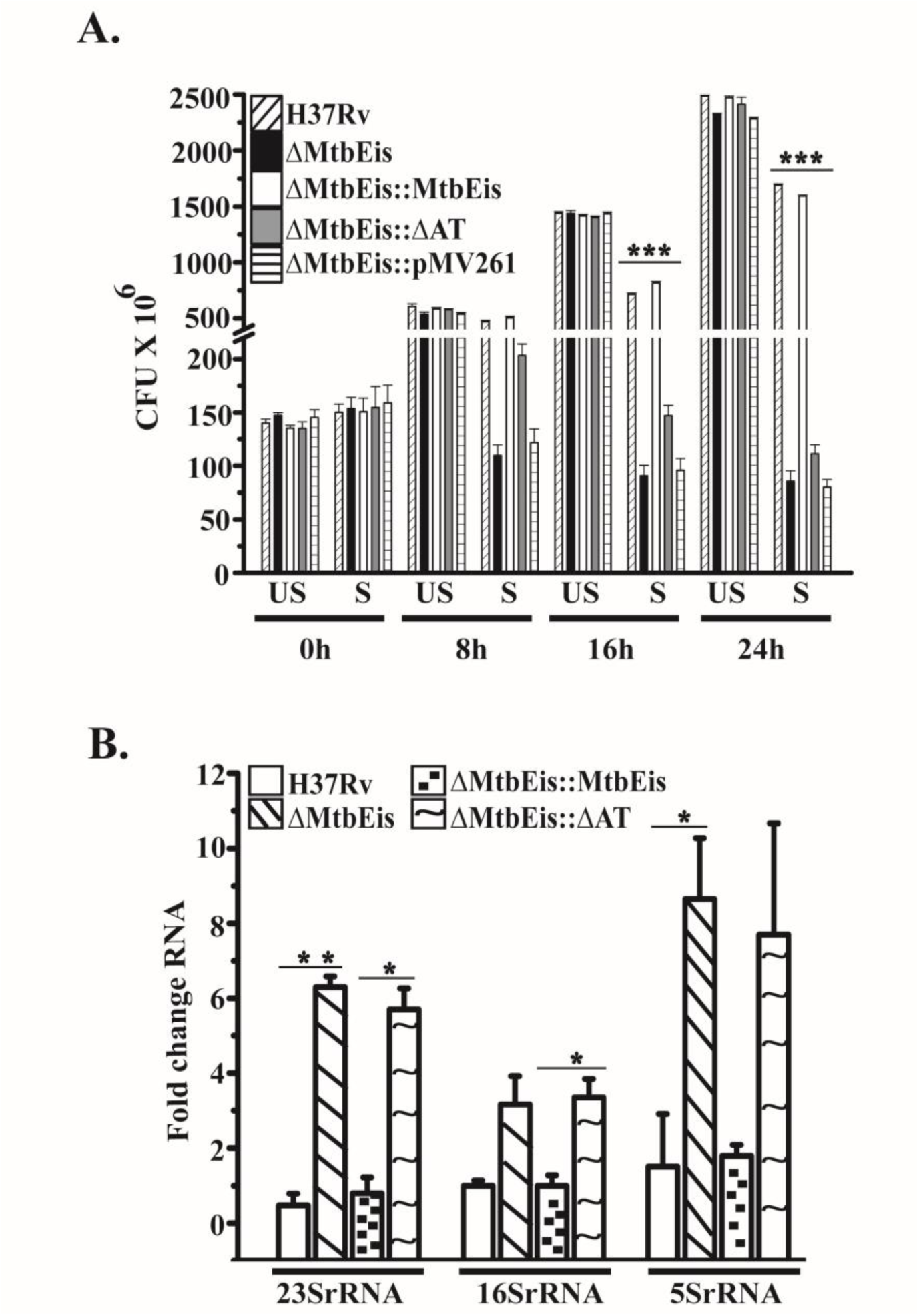
MtbEis mediated acetylation influences the survival of *M. tuberculosis* by inhibiting synthesis of rRNAs under starvation conditions. **Panel A**. The starvation survival of wild type H37Rv, ΔMtbEis and complemented strains were determined by CFU assay at 0 h, 8 h, 16 h and 24 h under unstarved (US) and starved (S) conditions. The average CFU counts obtained from three independent experiments were plotted (n=3). **Panel B.** The RNA was isolated from starved cultures of wild type H37Rv, ΔMtbEis and complemented strains. The levels of rRNAs were determined using RT-qPCR. The error bars depict mean standard deviation calculated from two independent experiments (n=2). The significance values were calculated by Student’s t-test analysis. * p-value ≤ 0.05, ** p-value ≤ 0.005, *** p-value ≤ 0.0005. **Figure 4-Source data 1** **Source data 1 file for Figure 4 includes an excel file of the compiled data used to generate image in Panel A**. **Figure 4-Source data 2** **Source data 2 files for Figure 4 include excel files containing raw and compiled data for RT-PCR analysis used to generate image in Panel B**.

**Fig. 5:**
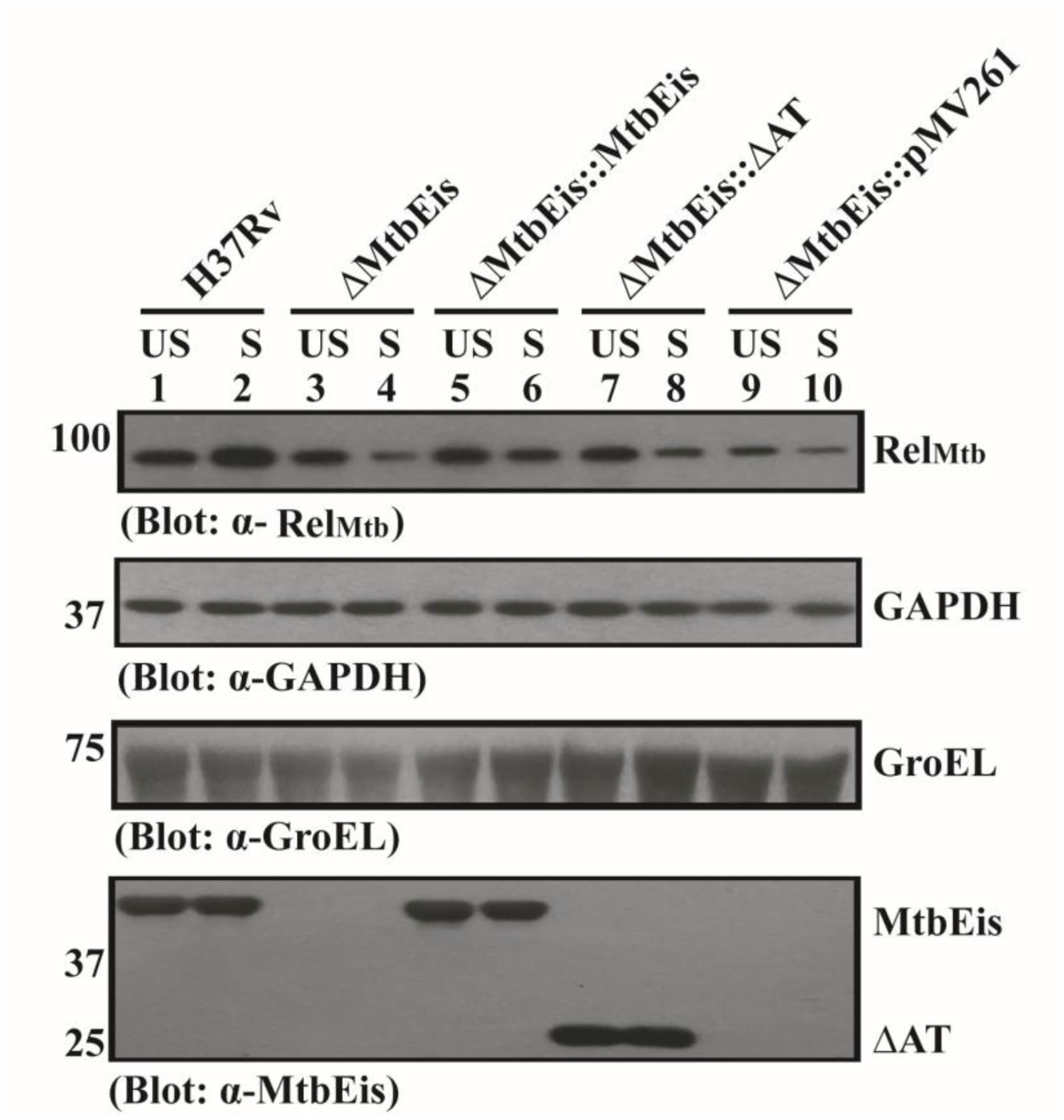
MtbEis mediated acetylation of Rel_Mtb_ regulates its expression under starvation conditions: Whole cell lysates prepared from unstarved (US) and starved (S) cultures of wild type H37Rv, ΔMtbEis and complemented strains were electrophoresed, transferred onto nitrocellulose membrane and probed with anti-Rel_Mtb_ antibody for the detection of Rel_Mtb_. MtbEis and ΔATEis proteins were detected using and anti-MtbEis antibody. The detection of GAPDH and GroEL using anti-GAPDH and anti-GroEL antibodies, respectively, served as loading control. The molecular weight markers and names of the proteins are shown on left and right side, respectively. The blots are representation of two independent experiments. **Figure 5-Source data 1** **Source data 1 files for Figure 5 include raw and labelled images for the blots shown in the figure 5**.

### Acetyltransferase domain of MtbEis regulates accumulation of (p)ppGpp under nutrient deprived conditions

Due to the reduced expression of Rel_Mtb_ upon deletion of MtbEis, it is important to validate the role of MtbEis in accumulation of (p)ppGpp during stringent conditions. Deletion of MtbEis resulted in abrogation of accumulation of (p)ppGpp which was restored on complementation with full length MtbEis (Fig.6, Lanes 1 vs 2 vs 3). However, ΔAT failed to restore the accumulation of (p)ppGpp (Fig.6, Lanes 1 vs 4). This result establishes the importance of MtbEis mediated acetylation of Rel_Mtb_ in regulating the accumulation of (p)ppGpp in Mtb.

## Discussion

Stringent response plays an important role in survival of bacteria under stress conditions. This response is mainly characterized by the down-regulation of stable RNAs, metabolism of carbohydrates, amino acids, phospholipids and upregulation of enzymes involved in the amino acid synthesis etc (20). Besides playing a role in metabolic slow down, stringent response has also been implicated in affecting virulence in Mtb (7). Virulence factors of Mtb are necessary for establishing successful infection, acquisition of available nutrients and to evade intracellular immune response by regulating stress response pathways (21). The differential expression of various virulence factors during activated stringent response in Mtb is required for the survival of mycobacterium (10). MtbEis is one of the virulence factor differentially regulated by Rel_Mtb_ and also implicated in drug resistance, autophagy and modulation of host immune response (10,12). However, the role of MtbEis in stringent response is not known.

Here, we have identified the interaction of MtbEis with Rel_Mtb_ and the enhanced accumulation of (p)ppGpp due to MtbEis mediated acetylation of Rel_Mtb_ at K513 residue. Our studies for the first time discover Rel_Mtb_ as an acetylation substrate of MtbEis. Although the acetylation mediated regulation of various proteins/enzymes has been documented (22), the regulation of Rel_Mtb_ by post translational modification is completely unknown. The findings reported here about enhanced accumulation of (p)ppGpp due to MtbEis mediated acetylation of Rel_Mtb_ leading to the metabolic adaptation of Mtb to nutritional stress underline a new mechanism regulating stringent response. The downregulation of Rel_Mtb_ expression with corresponding accumulation of (p)ppGpp in the absence of MtbEis confirms the regulation of stringent response by MtbEis mediated acetylation. The (p)ppGpp is a molecular alarmone that binds to the RNA polymerase and alters its promoter specificity so as to induce transcription of genes promoting survival of bacteria under stress conditions. Thus, the transcriptional and translational processes of cells are modulated to adjust the metabolism of cell according to the existing environmental challenges (23). The regulations of transcriptional and translational processes are both energy and time consuming for the cell. The post-translational modifications (PTMs) of proteins being more efficient mechanism is probably an alternative way for faster execution of regulations by diverting cellular resources away from unnecessary transcriptional and translational reactions for rapidly altering physiology of cells to assist bacterial survival under unfavorable conditions (22,24).

The reports on significance of acetylation of proteins affecting their functions such as modulation of enzymatic activities etc. are well documented. Some of the examples include alteration in the DNA binding ability of acetylated MtHU of *M. tuberculosis* (16); acetylation of MreB protein of *B. subtilis* leads to cell size alteration in concordance with the reduction in diameter of cell and restricted cell wall growth (25); decrease in activity of RNaseII due to enhanced acetylation of RNaseII of *E. coli* during slow growth conditions (26); adaptation of *M. smegmatis* in iron limiting conditions due to acetylation of MbtA (27); adaptation of Salmonella due to enhanced acetylation of metabolic enzymes when carbon source is altered (28) and regulation of DNA damage response during stationary phase by multiple acetylation of Ku protein in *M. smegmatis* (29). The concomitant escalation of acetylation modifications during the transition from actively growing bacterial cells to stationary phase depicts the tight regulation of physiological changes upon nutrient depleted conditions (30–32). Thus, the concept of acetylation as a regulatory process is irrefutable. Our results also highlight the role of MtbEis mediated acetylation in the survival of mycobacterium under nutrient limited conditions by altering the levels of rRNAs.

MtbEis is a well-known acetyltransferase responsible for acetylating aminoglycosides as well as proteins. Acetylation of aminoglycosides causes resistance against drugs used in second line treatment of Tuberculosis and acetylation of various proteins alters their function (12). The crucial role of MtbEis is evident when knock out strain of MtbEis exhibited compromised survival specifically under starved conditions which is restored by the full length MtbEis but not by the strain where N-terminus (1–161) is deleted. The N-terminus of MtbEis is known to harbor acetyltransferase domain (33), therefore, it is plausible that the acetylation activity of MtbEis is important for the execution of its role under stringent conditions. The findings of this study have been summarized in Fig 7.

**Fig. 6:**
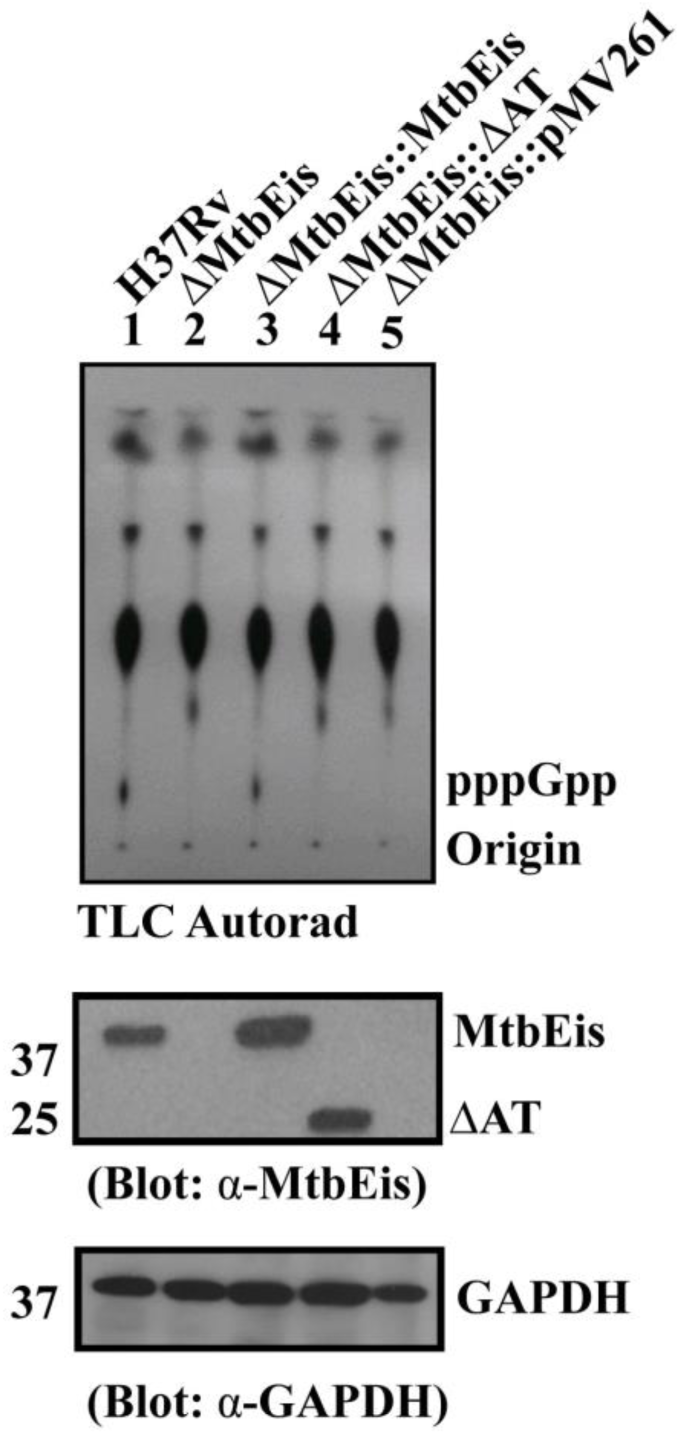
Acetyltransferase domain of MtbEis regulates accumulation of (p)ppGpp: The (p)ppGpp synthesis assay using whole cell lysates prepared from the starved H37Rv wild type, ΔMtbEis and complemented strains were performed and resolved on PEI–TLC plates followed by autoradiography. The blots are representation of two independent experiments. **Figure 6-Source data 1** **Source data 1 files for Figure 6 include raw and labelled images for the blots shown in the figure 6**.

**Fig. 7:**
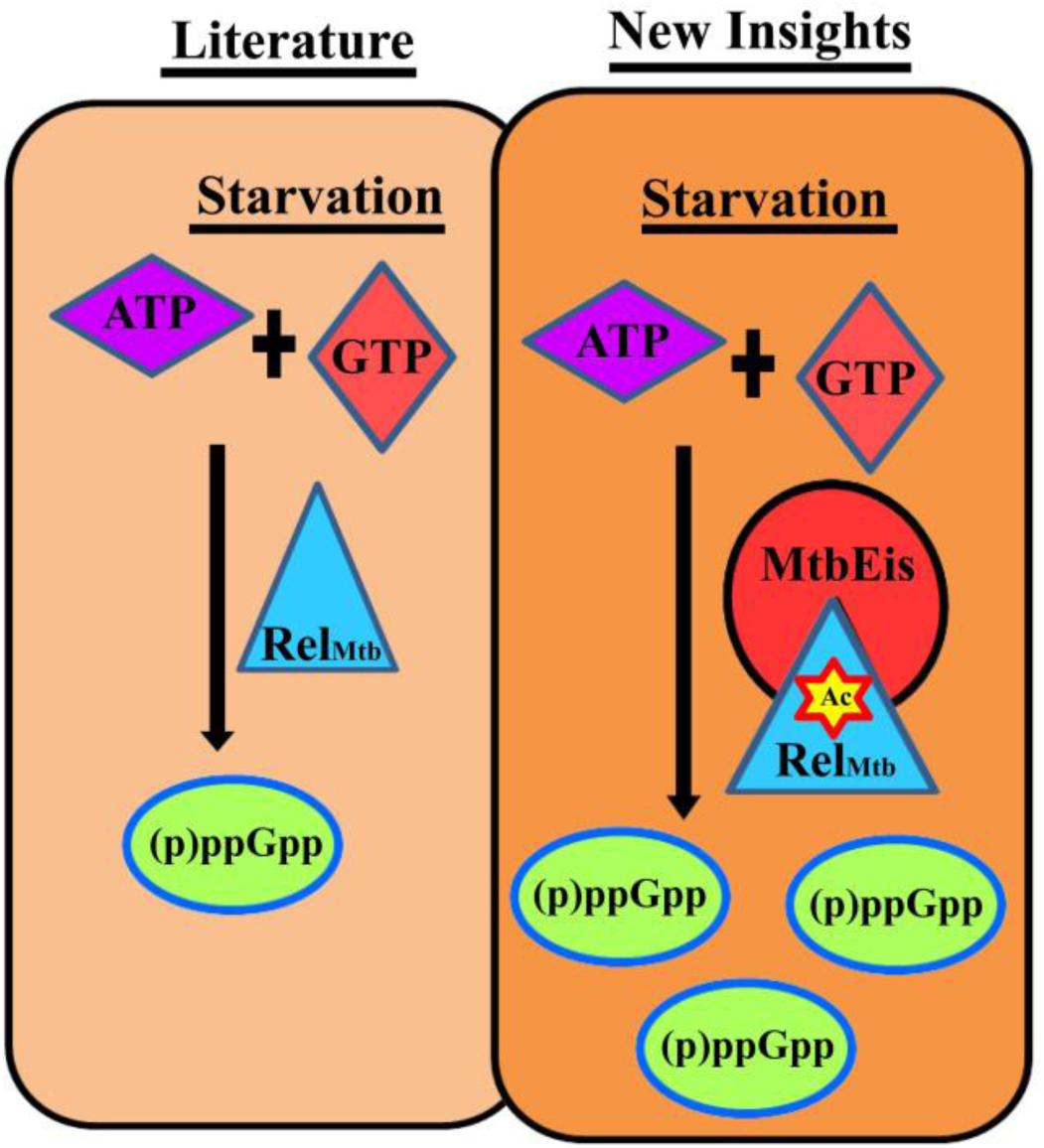
Model illustrating the impact of MtbEis mediated acetylation of Rel_Mtb_ in stringent response regulation: **(Left)** According to the information already available in existing literature, Rel_Mtb_ protein catalyzes the synthesis of (p)ppGpp from ATP and GTP substrates under starvation conditions. **(Right)** The novel insights drawn from our results assign a new function to MtbEis in stringent response. In sum, the interaction of Rel_Mtb_ with MtbEis results into its acetylation leading to enhanced synthesis of pppGpp under starvation conditions.

Also, our studies have hinted that the role of MtbEis in the survival is not apparent when there is a feast situation. There have been varying reports on the role of MtbEis in intracellular survival (12). However, in this study we have seen the crucial role of MtbEis only under scarcity of nutrients. Therefore, our studies suggest that it is important to review the growth conditions before assessing the function of MtbEis in the intracellular survival of mycobacterium.

In nut shell, these studies assign a novel role to MtbEis by regulating the stringent response through acetylation of Rel_Mtb_ and adding a new dimension about assessing the intracellular survival function of MtbEis. This study has also opened up the field for exploring novel post translational modifications regulating stringent response in mycobacterium.

## Materials and Methods

### Reagents

Restriction endonucleases were procured from New England Biolabs (NEB). Oligonucleotide primers were purchased from Bioserve. RNeasy^®^ Kit for RNA purification was obtained from QIAGEN. SuperScript^®^ III Platinum^®^ SYBR^®^ Green One-Step qRT-PCR Kit was bought from Invitrogen. GST and HRP Conjugated secondary antibodies for Western blotting were procured from Santa Cruz Biotechnology. Anti-MtbEis, anti-Rel_Mtb_ antibodies were raised in rabbit in house. The ɣ^32^P and ^14^C-Acetyl CoA was purchased from BRIT, India and Perkin Elmer, respectively. Lysing Matrix B was obtained from MP Biomedicals. All analytical and molecular biology grade chemicals were purchased from Merck.

### Bacterial strains, media and growth conditions

The DH5α and BL21(DE3)pLysS were grown in LB. The required concentrations of Kanamycin (50 µg/ml), Chloramphenicol (34 µg/ml), Ampicillin (100 µg/ml) and Hygromycin (200 µg/ml) antibiotics were added as required for *E. coli* strains. *M. tuberculosis* H37Rv was grown at 37^ᵒ^C either in Middlebrook 7H9 media (Difco) along with 10% OADC (BD), 0.2% glycerol, 0.05% Tween-80 or on 7H11 agar containing 0.2% glycerol and 10% OADC. Kanamycin (20µg/ml) and Hygromycin (50µg/ml) were used for the growth of *M. tuberculosis* as required.

### Purification of GST tagged proteins

GST-Rel_Mtb_ protein purification was done according to the protocol published by Gozani lab in 2005 (34) . GST-Rel_Mtb_ protein was induced in BL21(DE3)pLysS cells using 2mM IPTG for 6.0 h at 28^ᵒ^C. The harvested cells were sonicated in lysis buffer containing 50 mM Tris (pH 7.5), 150 mM NaCl and 0.05% NP-40. The supernatant fraction of cleared cell lysate was subjected to end-to-end binding with glutathione agarose beads mixture for 12 h at 4^ᵒ^C. The beads were washed thrice with lysis buffer followed by another washing with elution buffer containing 50 mM Tris (pH 8.0) and 300 mM NaCl. The protein was eluted in 1.0 ml of elution buffer supplemented with 10 mM reduced glutathione. The eluted protein was dialyzed against elution buffer without glutathione and quantified using Bradford assay. GST protein was purified from pET41a vector which is having both His tag and GST tag.

### GST-Pull Down Assay

Equimolar concentrations of purified recombinant GST, GST- Rel_Mtb_ and MtbEis proteins were allowed to bind with glutathione agarose beads in pull down buffer having 20 mM Tris (pH 8.0), 150 mM NaCl, 0.5% NP-40, 10% glycerol and 1mM MgCl_2_. The eluted fractions obtained by boiling beads in 1X Laemmli buffer were electrophoresed onto 10% SDS gel, Western transferred and immunoblotted using anti-HIS, anti-MtbEis and anti-GST antibodies.

### Preparation of whole cell lysates of *M. tuberculosis*

To prepare whole cell lysates of all mycobacterial strains, fully grown harvested cultures were washed twice with 1X PBS and resuspended in 1X PBS containing protease inhibitor cocktail. The cells were lysed using lysing matrix B by bead beater at 6 m/s for 30 s for 12 cycles with intermittent incubation on ice for 5 min each followed by centrifugation at 13,000 rpm for 10 min. The 0.22 µ filtered supernatant was quantified using Bradford reagent. For RNA preparation, cell suspension was subjected to only four cycles of bead beating.

### Co-Immunoprecipitation of endogenous Rel_Mtb_ and MtbEis

The varying concentration of Rel_Mtb_ antibody along with Protein A beads was used for immunoprecipitating MtbEis from Mtb lysates by mixing at 4^°^C for 12 h. The harvested beads were washed thrice with 1X PBS followed by boiling in 1X Laemmli buffer. The supernatant was electrophoresed on 10% SDS-PAGE and after Western blotting proteins were detected using anti-Rel_Mtb_ and anti-MtbEis antibodies.

### Purification of His-MtbEis and His-Rel_Mtb_ proteins

Purification of His-MtbEis was done as described previously (35). For Rel_Mtb_ protein, 2.0 mM IPTG was used to induce the expression in BL21(DE3)pLysS cells for 6.0 h at 28^°^C. The harvested cells were washed with 1X PBS and sonicated in phosphate buffer (pH 8.0) containing 50 mM NaH_2_PO_4_, 300 mM NaCl, 10 mM Imidazole and protease inhibitor cocktail mix. The Ni-NTA column loaded with cleared lysate was washed with phosphate buffer containing 10 mM, 20 mM, 30 mM, 40 mM, 50 mM and 100 mM Imidazole. The protein was eluted in phosphate buffer (pH 8.0) containing 250 mM Imidazole. The eluted fractions were dialyzed extensively against 20 mM sodium phosphate buffer (pH 8.0) containing 300 mM NaCl. The yield of the purified protein was 0.5 mg/liter.

### Guanosine-pentaphosphate synthesis assay

The pppGpp synthesis assay was performed with slight modification of previously described method (6). Briefly, the pppGpp synthesis was carried out at 37^ᵒ^C for 30 min in 10 µL reaction mixture containing 20 mM phosphate buffer (pH 8.0) consisting of 300 mM NaCl, 5.0 mM MgCl_2_, [ɣ^32^P]ATP (1.0 µCi/µmol), 5.0 mM GTP and 2.5 µM of desired proteins (Rel_Mtb_/MtbEis/Rel_Mtb_K513A). The reaction was stopped by addition of 2.0 µl of 6N Formic acid followed by centrifugation at 13,000 rpm for 10 min at 4^ᵒ^C. The 5.0 µl of the supernatant was spotted on TLC PEI Cellulose plate and allowed to resolve in 1.5 M KH_2_PO_4_ (pH 3.4) followed by air drying and autoradiography.

### *In vitro* Acetyltransferase assay

The acetylation assay was performed at 37^ᵒ^C for 2 h in reaction mixture containing 0.2 µg of His-MtbEis and 5.0 µg of His-Rel_Mtb_/Rel_Mtb_ (K513A), 100 µM ^14^C-Acetyl CoA (63 mCi/mmol), 10 % (v/v) glycerol, 1.0 mM DTT, 1.0 mM Trichostatin A, 0.1 mM EDTA, and 50 mM Tris-Cl (pH 8.0). The reaction was arrested by boiling in 1X Laemmli buffer, electrophoresed by SDS-PAGE and transferred onto nitrocellulose membrane. The incorporation of ^14^C was analysed using autoradiography. The anti-His antibody was used for probing the proteins used in the assay.

### LC-MS/MS detection of residues of acetylation

Following *in vitro* acetylation assay, the excised corresponding bands of acetylated and unacetylated Rel_Mtb_ proteins obtained from SDS-PAGE were sent for LC-MS/MS analysis to Taplin Biological Mass Spectrometry facility at Harvard Medical School (http://taplin.med.harvard.edu/). After data analysis the graphs were plotted using Origin^®^ 9.1 software.

### Construction of RvEis knockout

The left homology sequence (LHS) and right homology sequence (RHS) of MtbEis were amplified using specific primer pairs enlisted in Table S2. Restriction digestion of amplified fragments (LHS & RHS), vectors p0004S and pENTR with pfIMI resulted in ends compatible for ligating LHS, RHS, hyg cassette, λcos sites to generate the allelic exchange substrate (AES) in a single step. AES was linearized by restriction digestion using EcoRV. To create *eis* gene replacement mutants, H37Rv transformed with pNit-ET in log phase was treated with 5µM isovaleronitrile followed by 0.2 M glycine and allowed to grow till O.D_600_ 0.8. The harvested cells were washed for five times with 10% glycerol and electroporated with 500 ng of linear AES. The transformed colonies were screened for gene replacement by PCR. The putative positive clones were confirmed by RT PCR using gene specific primers and Western blotting using α-Eis antibody. For complementation of MtbEis knockout, pMV261 vector carrying full length MtbEis (36) and MtbEis lacking acetyltransferase domain (ΔAT) were electrotansformed in pNit-ET cured ΔMtbEis cells.

### Starvation survival assay

All mycobacterial strains grown till exponential phase were split in two bottles. Starvation was induced for 24 h in one set by washing harvested cells twice with 1X PBS followed by suspension in TBST and incubation at 37^ᵒ^C. The cells in other culture bottle were allowed to grow. Samples were taken at different time points for cfu assay from respective starved and unstarved cultures. The graphs were plotted using Origin^®^ 9.1 software.

### Mtb RNA isolation

All harvested Mycobacterial cultures were washed with TE buffer (pH 8.0) and resuspended in 1.0 ml TRIzol reagent. The cells were lysed as described above. After Phenol chloroform extraction, RNeasy Protect Bacteria Mini Kit was used for the extraction of RNA as per the manufacture’s protocol. DNAase I treatment was done to remove any residual genomic DNA.

### Quantitative Real-Time PCR

Quantitative Real-Time PCR was performed using one step RT-PCR kit with suitable primers (200nM) in the Applied Biosystem’s one step plus PCR thermocycler. Mtb *groEL2* expression was used as a control to normalize each sample in all experiments. Fold difference was calculated using ΔΔC_T_. Each experiment was repeated twice with independent RNA samples to calculate average fold differences and standard deviation.

### Guanosine-pentaphosphate synthesis assay with Mtb lysates

Exponential grown all Mycobacterial strains were subjected to complete starvation in TBST at 37^ᵒ^C for 24 h. The whole cell lysates prepared as mentioned above were concentrated in pierce protein concentrators (MWCO 3kDa) and quantified using Bradford reagent. The assay was set up with 30 µg of lysates in 20 µl reaction mixture as mentioned above. Only 10 µl was used for analysis.

### Statistical significance

The graphs were plotted using Origin^®^ 9.1 software. Absolute values are average of at least two/three biological replicates ± standard deviations and are representative of at least two independent experiments. The statistical significance was calculated by GraphPad software using Student’s t-test for the starvation Survival assay and RT-qPCR experiments.

### Data availability

The main data generated in this study is available in the paper and its supplementary figures. Raw data files are available from the corresponding author upon appropriate request. Source data files are provided with the paper.

## Acknowledgements

This work was funded by Council of Scientific and Industrial Research (CSIR, grant number MLP050), New Delhi, India. The funding source had no role in the design of the study, data analysis, decision to publish or the writing of the manuscript. Fellowship to NS and VB was from Department of Biotechnology (DBT), New Delhi and CSIR, New Delhi, India respectively. Dr. Eric Rubin, from Howard School of Public Health, USA is acknowledged for the gift of pNit-ET plasmid. Dr. Vinay Nandicoori, Director, Centre for Cellular and Molecular Microbiology is acknowledged for the kind gifts of p0004S and pENTR plasmids. Dr. Ritesh Rajesh Sevalkar from Dr. Dibyendu Sarkar’s group at CSIR-IMTECH is acknowledged for helpful discussions for MtbEis knockout generation. We also thank Dr. Manoj Raje, CSIR-IMTech, Chandigarh and Prof Chaaya Iyengar Raje, NIPER, Mohali for providing Mtb-GAPDH antibody.

## Author Contributions

C.S. conceived and designed the studies and procured funding. All the experiments were performed by N.S. V.B performed site directed mutagenesis, recombinant protein preparations and assisted in starvation survival assays. N.S. and C.S. analyzed the data and prepared the manuscript.

## Supplementary Information

The supplementary data for this article is available online. The Source data files for supplementary data are provided with the paper.

## Materials Availability Statement

The newly created materials described in this study can be accessed upon request. All the data sets have been deposited in the institutional repository.

## Supplementary data

**Fig. S1:**
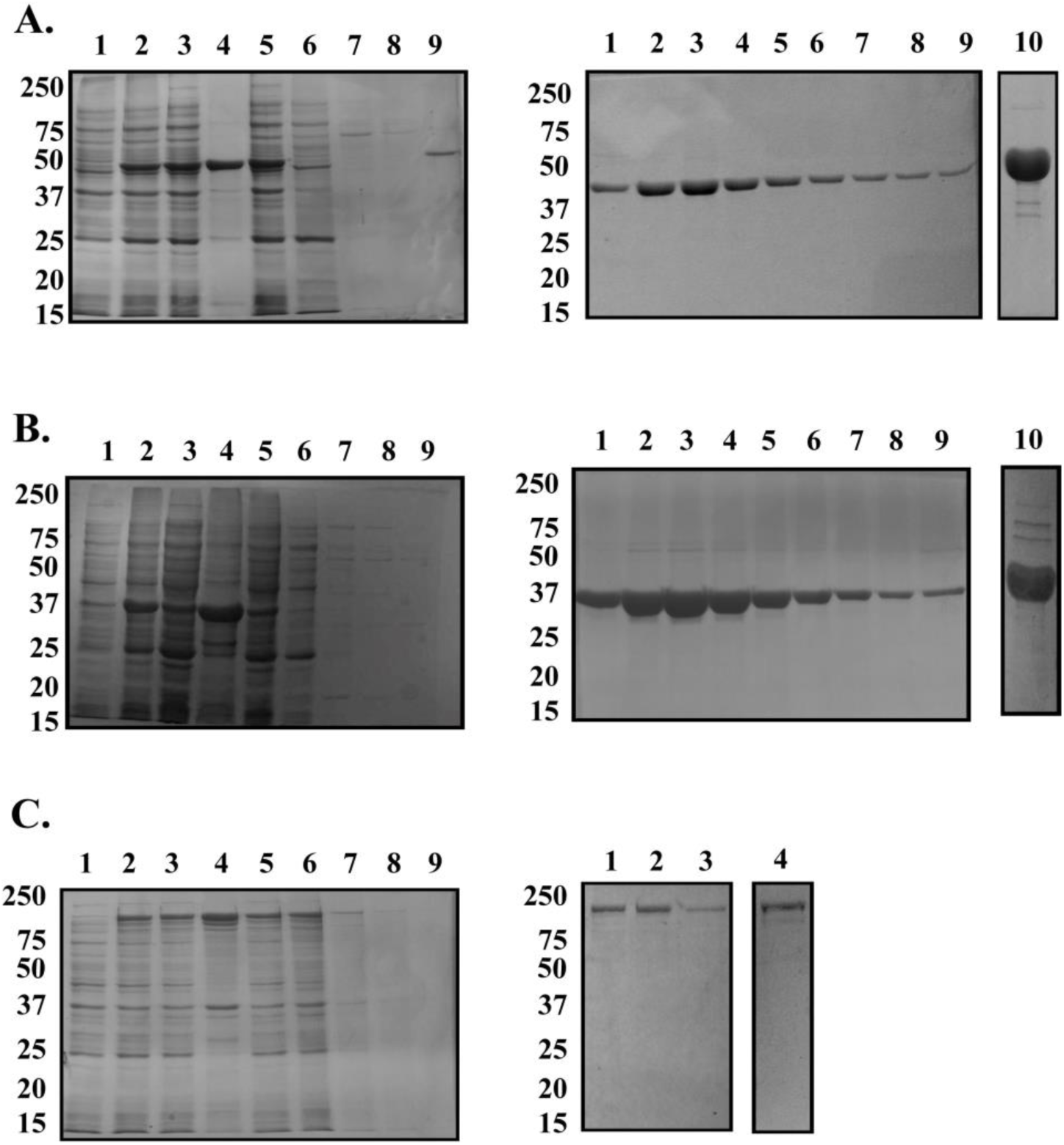
Purification of His-MtbEis, GST and GST-Rel_Mtb_ proteins. **A: Purification of His-MtbEis. Left Panel:** Lane 1: Uninduced; Lane 2: Induced; Lane 3: Load; Lane 4: Pellet; Lane 5: Flow through; Lanes 6-9: Washes with increasing concentration of Imidazole. **Right Panel:** Lanes 1-9: Elution fractions of His-MtbEis protein with buffer containing 250 mM Imidazole; Lane 10: Concentrated MtbEis protein after dialysis. **B: Purification of GST. Left Panel:** Lane 1: Uninduced; Lane 2: Induced; Lane 3: Load; Lane 4: Pellet; Lane 5: Flow through; Lanes 6-9: Washes with increasing concentration of Imidazole. **Right Panel:** Lanes 1-9: Elution fractions of GST protein with buffer containing 250 mM Imidazole; Lane 10: Concentrated GST protein after dialysis. **C: Purification of and GST-Rel**_Mtb_**. Left Panel:** Lane 1: Uninduced; Lane 2: Induced; Lane 3: Load; Lane 4: Pellet; Lane 5: Flow through; Lanes 6-8: Washes with lysis buffer; Lane 9: Wash with elution buffer without glutathione**. Right Panel:** Lanes 1-3: Elution fractions of GST-Rel_Mtb_ protein with buffer containing 10 mM reduced glutathione; Lane 4: Concentrated GST-Rel_Mtb_ protein after dialysis. Molecular weight markers for all the panels are shown on left side.

**Fig. S2:**
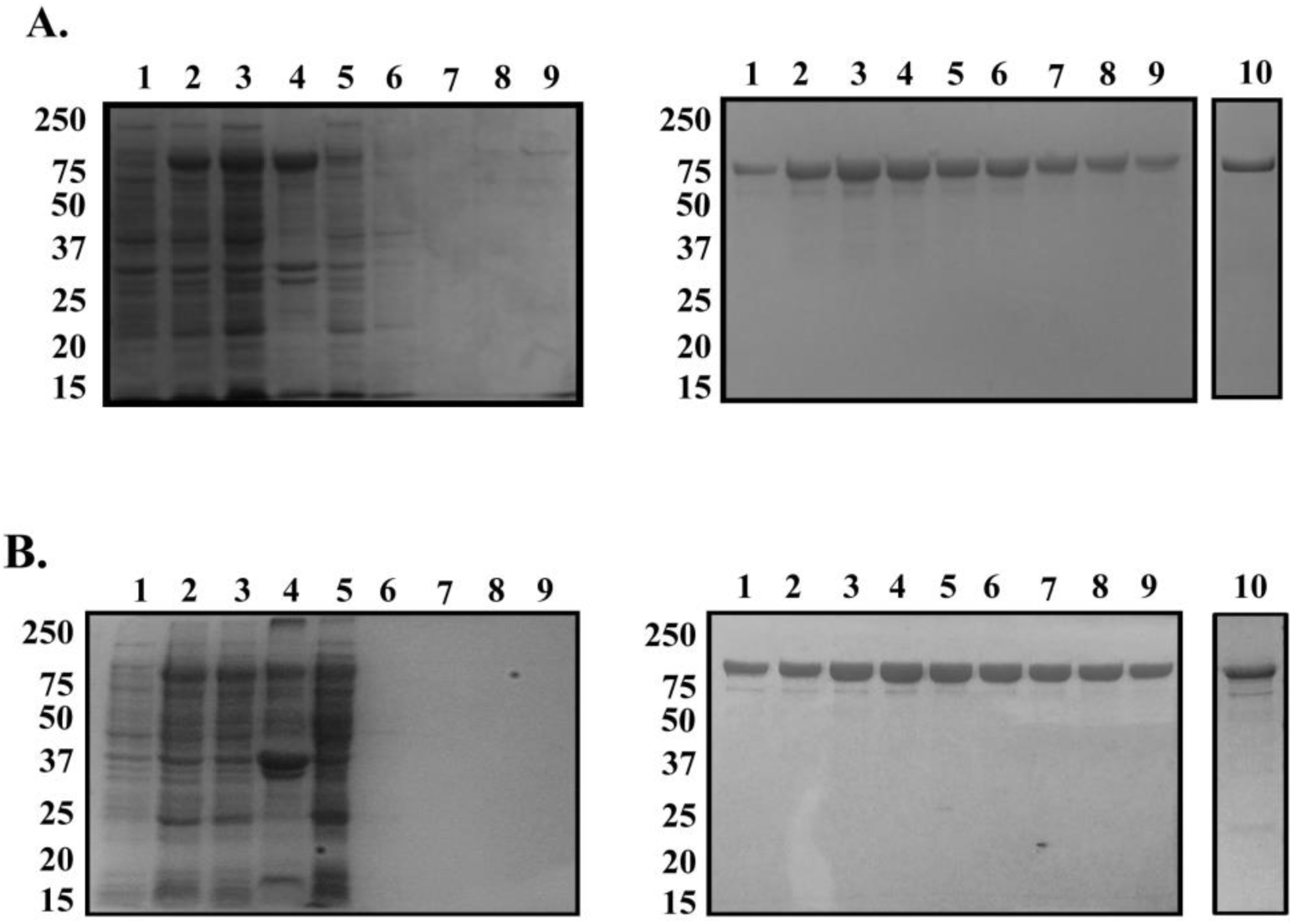
Purification of His-Rel_Mtb_ and His-Rel_Mtb_(K513A) proteins. **A: Purification of His-Rel**_Mtb_**. Left Panel:** Lane 1: Uninduced; Lane 2: Induced; Lane 3: Load; Lane 4: Pellet; Lane 5: Flow through; Lanes 6-9: Washes with increasing concentration of Imidazole**. Right Panel:** Lanes 1-9: Elution fractions of His-Rel_Mtb_ protein with buffer containing 250 mM Imidazole; Lane 10: Concentrated Rel_Mtb_ protein after dialysis. **B: Purification of His-Rel**_Mtb_**(K513A). Left Panel:** Lane 1: Uninduced; Lane 2: Induced; Lane 3: Load; Lane 4: Pellet; Lane 5: Flow through; Lanes 6-9: Washes with increasing concentration of Imidazole**. Right Panel:** Lanes 1-9: Elution fractions of His- Rel_Mtb_(K513A) protein with buffer containing 250 mM Imidazole; Lane 10: Concentrated Rel_Mtb_(K513A) protein after dialysis. Molecular weight markers for all the panels are shown on left side.

**Fig. S3:**
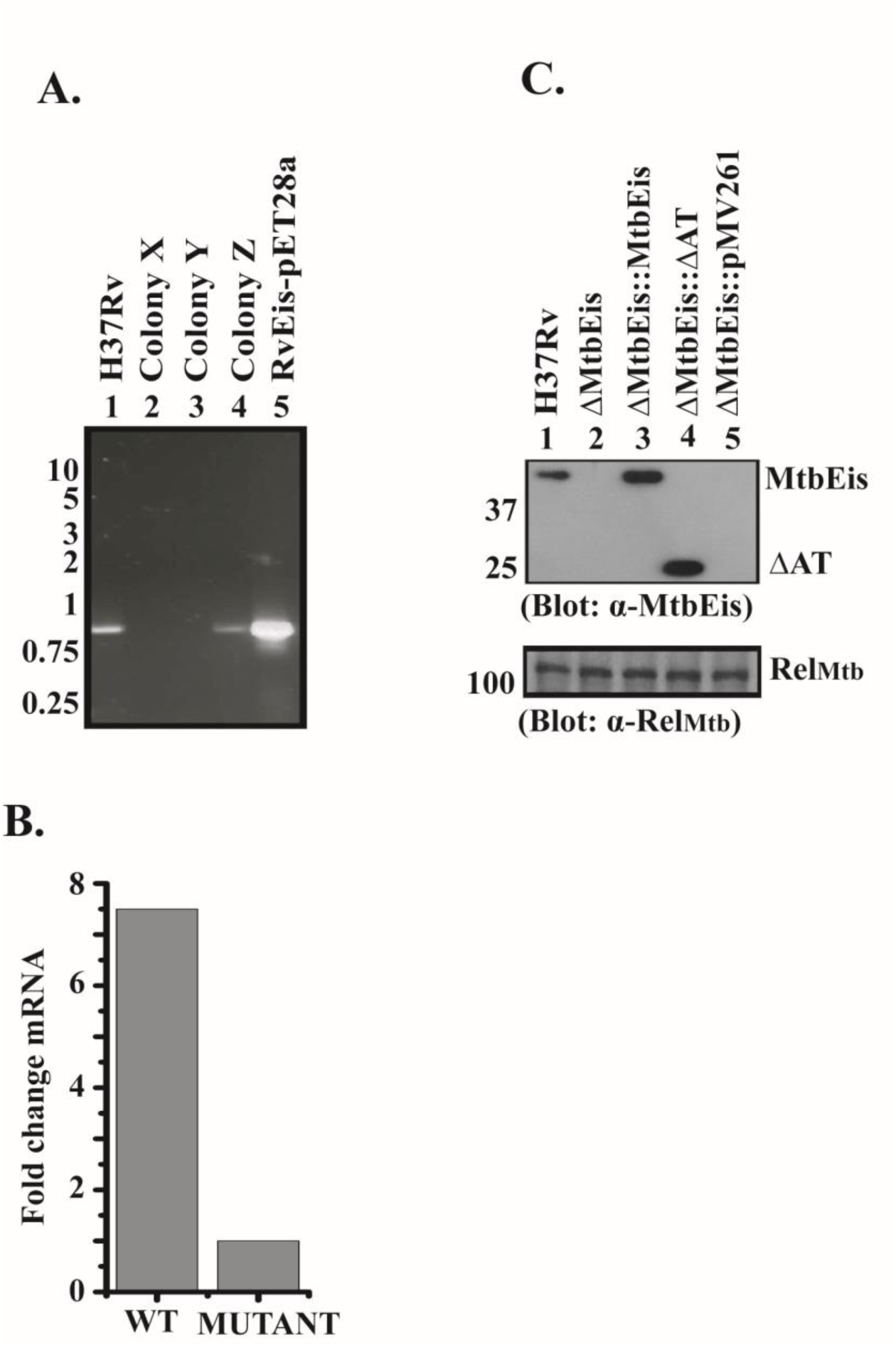
Screening, confirmation and complementation of ΔMtbEis. **A**: Screening of MtbEis gene replacement mutants by PCR using internal primers specific to MtbEis. Molecular weight markers (kb) are shown on left side **B.** Confirmation of MtbEis knockout creation by q-RT PCR. **C:** Confirmation of MtbEis knockout generation by western blotting (Lane 2) and complementation of ΔMtbEis with full length MtbEis (Lane 3) and MtbEis lacking acetyltransferase domain (Lane 4). Molecular weight markers are shown on left side.

## Supplementary tables

**Table S1:**
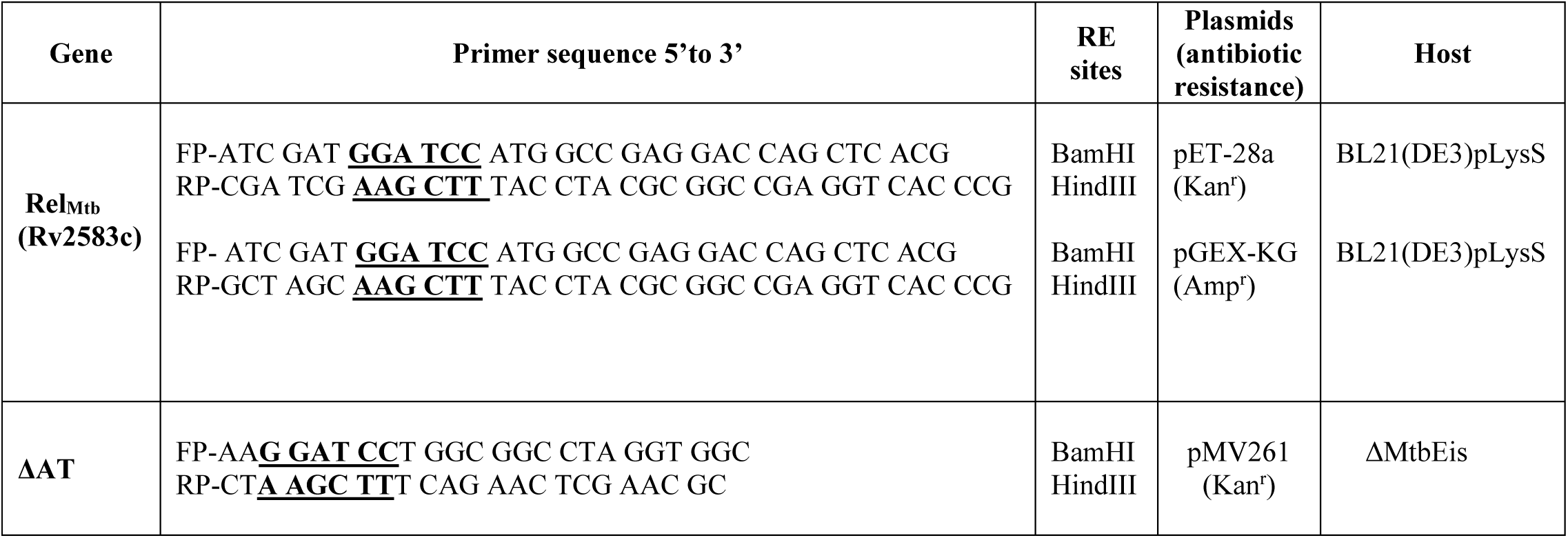
Genes, Primers, plasmids and bacterial strains used in the study.

**Table S2:**
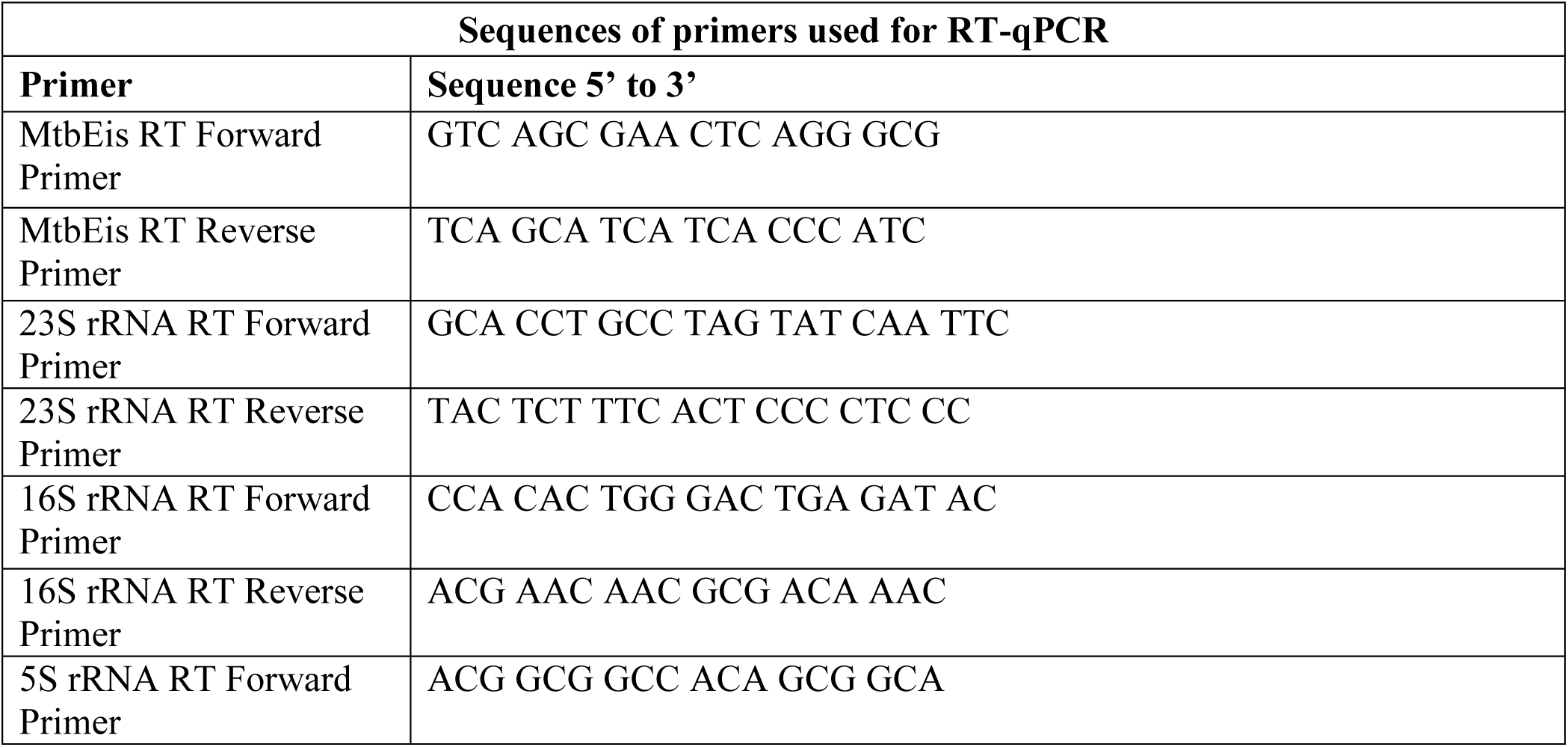
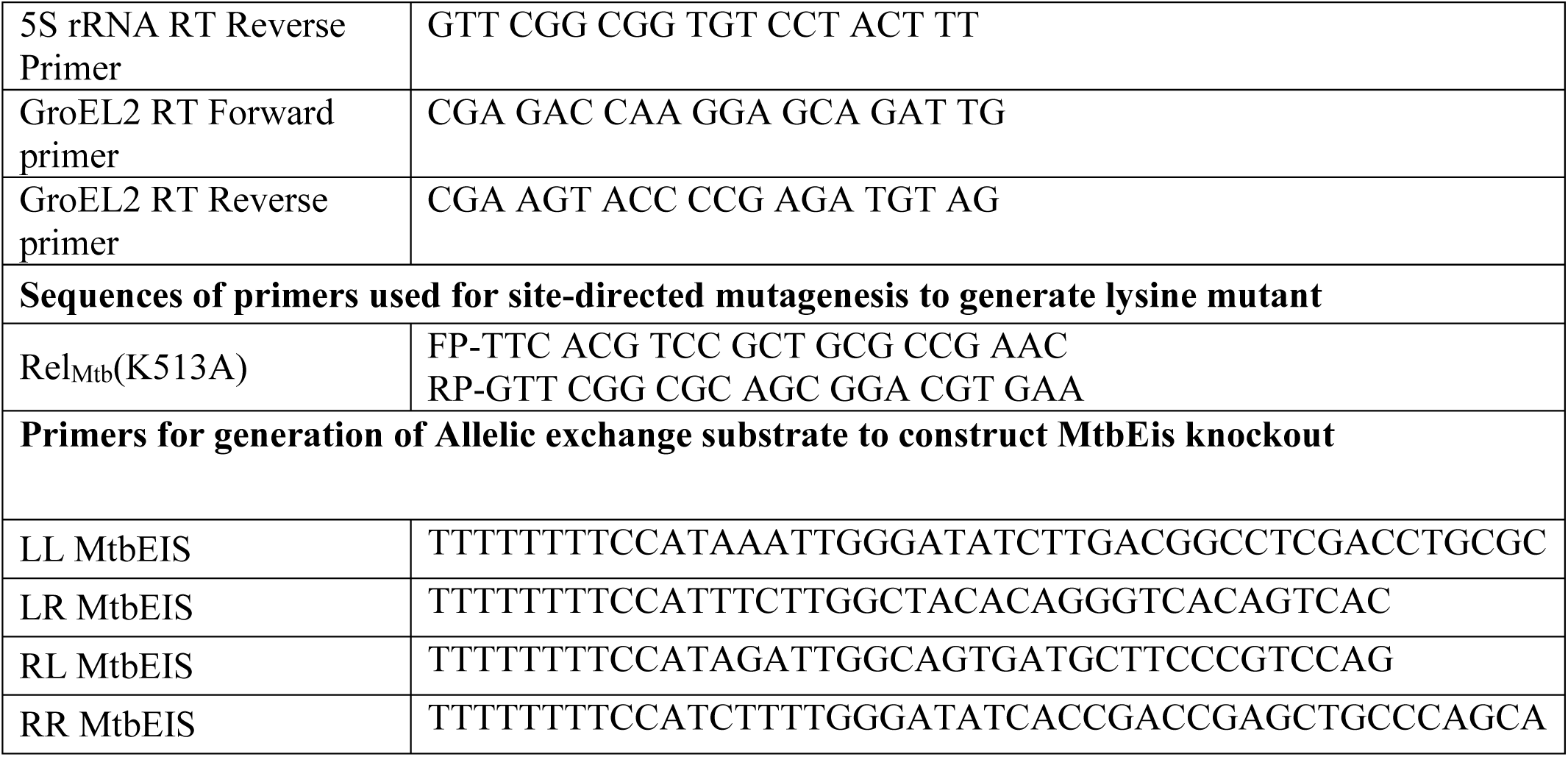
Primers used for RT-qPCR, SDM and AES generation.

